# Predicting the Impact of Dialyzer Choice and Binder Dialysate Flow Rate on Bilirubin Removal

**DOI:** 10.1101/2024.09.27.615470

**Authors:** Alexander Novokhodko, Nanye Du, Shaohang Hao, Ziyuan Wang, Zhiquan Shu, Suhail Ahmad, Dayong Gao

## Abstract

Liver failure is the 12^th^ leading cause of death worldwide. Protein bound toxins such as bilirubin are responsible for many complications of the disease. Binder dialysis systems use albumin dialysate and detoxifying sorbent columns to remove these toxins. Systems like the Molecular Adsorbent Recirculating System (MARS) and BioLogic-DT have existed since the 1990s, but survival benefit in randomized controlled trials have not been consistent. Thus, a new generation of binder dialysis systems, including Open Albumin Dialysis (OPAL) and the Advanced Multi-Organ Replacement System (AMOR) are being developed. Optimal conditions for binder dialysis have not been established. We developed and validated a computational model of bound solute dialysis using established thermodynamic theories. Our objective is to improve AMOR therapy. We confirmed our model’s validity by predicting the impact of changing between two benchtop dialysis setups using different polysulfone dialyzers (F3 and F6HPS). We then applied it to predict the impact of varying dialysate flow rate on toxin removal. We found that bilirubin removal is independent of dialysate flow rate within the clinically relevant range (20 mL/min – 800 mL/min), matching our model’s predictions. At very low dialysate flow rates (2 mL/min), bilirubin removal declines, deviating from the thermodynamic model. This model may be useful to achieving optimal clinical outcomes by setting optimal dialyzer and flow rate conditions. Further improvement is possible by accounting for toxin adsorption onto the dialyzer membrane.

## 1 Introduction

In 2021, Liver Failure was the 12^th^ leading cause of death globally (Naghavi et al., 2024). The liver performs a variety of critical functions, including toxin removal, drug and food metabolism, protein synthesis including synthesis of clotting factors and immunoproteins, nutrient storage, and other functions. The buildup of toxins during liver failure drives hepatic coma, liver cell death, and failure of other organ systems. Bilirubin is a commonly studied hepatic toxin. Bilirubin accumulation is visible in the skin as jaundice. Excess bilirubin contributes to the pathology of liver failure, for example by damaging white matter (Lakovic et al., 2014), which may contribute to hepatic coma. It is also a commonly accepted marker for other albumin-bound toxins.

Bilirubin cannot be removed by traditional dialysis because it is a hydrophobic molecule that is bound to albumin in human blood (Kalakonda et al., 2024). Adding a binder such as albumin or charcoal to the dialysate allows bilirubin removal (Patzer, 2006). Computational modeling of albumin dialysis was first done in the seminal work of Patzer and colleagues, who described single pass albumin dialysis (SPAD) in three publications (Patzer, 2006; Patzer et al., 2006; Patzer and Bane, 2003). Today, the only FDA approved albumin dialysis system is the Molecular Adsorbent Recirculating System (MARS) (Padmanabhan et al., 2019), which recirculates albumin instead of discarding it after a single pass. Four groups have published models of closed loop mode albumin dialysis.

The first is by Magosso and colleagues (Magosso et al., 2006). This model was designed using clinical data to model MARS sessions. It accounts for ultrafiltration and diffusion. However, it assumes that the dialyzer blood and dialysate compartments and charcoal and resin columns of MARS are well mixed compartments, without modeling concentration gradients. The model has not been validated against data beyond what was used to fit its parameters.

The second is by Annesini and colleagues (Annesini et al., 2009, 2011, 2014). This model applies chemical engineering techniques. Unfortunately, it cannot be applied to bilirubin because it assumes albumin concentration is far greater than toxin concentration. As shown in Table 1, this is not always the case for hyperbilirubinemia in liver failure.

**Table 1:** Albumin and bilirubin concentrations in liver failure can be of the same order of magnitude. Values are minima and maxima.

| Substance | Molecular Weight (Da) | Average Concentration in Liver Failure (g/L) | Average Concentration in Liver Failure (μM) |
| --- | --- | --- | --- |
| Human Serum Albumin | 66,438 (Fanali et al., 2012) | 9.6 - 56.2 (Bai et al., 2019) | 144 - 846 |
| Bilirubin | 584.7 ("PubChem Compound Summary for CID 5280352, Bilirubin," 2022) | 0.00111 - 0.473 | 1.9 - 809.8 (Bai et al., 2019) |

The third is by Pei and colleagues, who extended the work of Patzer to include recirculation and the effect of local ultrafiltration (Pei et al., 2013, 2014b, 2014a, 2015). Unfortunately, when we replicated their model, we were unable to reproduce their results. They reported results for three initial blood bilirubin concentrations: 14.7, 17.7 and 21.4 mg/dL. Reported modeled final blood bilirubin concentrations from Pei and colleagues are 7.7, 9.0 and 10.8 mg/dL. In our attempt to reproduce their model with their parameters, we got final bilirubin concentrations of 10.36, 11.08, 11.76 mg/dL.

Their code is not publicly available, and parts of their model, such as the equation for ultrafiltration rate in the Peclet Number formula, are not reported, making it difficult to identify why their data could not be reproduced. In our attempt to replicate the model we used Villarroel’s definition of Peclet Number (Villarroel et al., 1977).

The last effort was by Schiesser (“Liver Support Systems,” 2016). This model couples a 2 ordinary differential equation (ODE) patient model with an 8 partial differential equation (PDE) model for the extracorporeal device. It has not been validated against experimental data.

MARS has not shown consistent survival improvements in randomized controlled trials (Bañares et al., 2013; Saliba et al., 2013). Today, a new generation of binder dialysis systems are being developed. These include the Advanced Multi-Organ Replacement System (AMOR) (Ahmad et al., 2024), Open Albumin Dialysis (OPAL) (Stange et al., 2018), and High Efficiency MARS (HE-MARS) (Marangoni et al., 2014). Our goal is to optimize AMOR using this algorithm, but any binder dialysis system could be modeled in this way.

## 2 Methods

### 2.1 Model Description

The model was constructed similarly to that of Pei and colleagues (Pei et al., 2013, 2014b, 2014a, 2015). Blood and dialysate were assumed to be two well-mixed compartments containing albumin and the toxin of interest. Toxin removal was assumed to happen by diffusion and convection across a dialyzer membrane with blood and dialysate flowing counter-current. The removal of toxin across the dialyzer was solved at each time step by solving a spatial model of concentrations, pressures, and flow rates. This was used to update the blood and dialysate concentrations. The Peclet number was defined following previous work (Villarroel et al., 1977). Unlike the model of Pei and colleagues, a piecewise function was used to account for the change in the sign of convective flux across the dialysis membrane during backfiltration.

### 2.2 Variable Names

*P*_*b*_→blood pressure, *P*_*d*_ →dialysate pressure, *z* → distance along dialyzer fiber from blood inlet and dialysate outlet. µ_*b*_→viscosity of blood. µ_*d*_ →viscosity of dialysate. *Q*_*b*_→blood flow rate. *Q*_*d*_ →dialysate flow rate. *r*_*i*_ →inner dialysate fiber radius. *r*_*o*_ →outer dialysate fiber radius. *R*_*m*_ →inner radius of dialyzer shell (constrains the dialysate space outside the fibers). *n* →number of fibers. *L*_*p*_ →Membrane hydraulic permeability. *C*_*stlb*_→total blood bilirubin concentration. *C*_*stld*_ →total dialysate bilirubin concentration. *C*_*sb*_→free blood bilirubin concentration. *C*_*sd*_ →free dialysate bilirubin concentration. *C*_*atlb*_→total blood albumin. *C*_*atld*_ →total dialysate albumin. *K*_*free*_ *A* → a coefficient representing the diffusive transport of unbound bilirubin across the entire area of the dialyzer membrane. *K*_*B*_ →Albumin-bilirubin binding constant, *J*_*v*_ →local ultrafiltration flux. *f* → function of Peclet Number. σ→reflection coefficient of the membrane. *L* → dialyzer length. *pe* → Peclet Number, a dimensionless quantity relating convection and diffusion. *V*_*b*_→blood reservoir volume. *V*_*d*_ →dialysate reservoir volume.

### 2.3 Boundary Conditions

*P*_*d*_(*z* = 0) = 0 (dialysate outlet pressure), *C*_*stlb*_(*z* = 0, *t* = 0) = *C*_*stlb,in*_ (input blood side bilirubin). *C*_*stld*_(*z = L, t* = 0) = 0 (dialysate initially has no bilirubin). *Q*_*b*_(*z = L*) *= Q*_*b*_(*z =* 0) (no net ultrafiltration). *Q*_*d*_(*z = L*) *= Q*_*d*_(*z =* 0) (no net ultrafiltration). *Q*_*b*_(*z =* 0) *= Q*_*b,in*_ (blood inlet flow rate). *Q*_*d*_(*z = L*) *= Q*_*d,in*_ (dialysate inlet flow rate).

### 2.4 Equations

Change in blood pressure with respect to

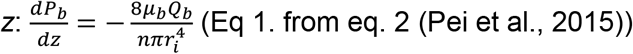

Change in dialysate pressure with respect to

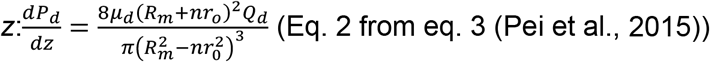

Change in blood flow rate with respect to

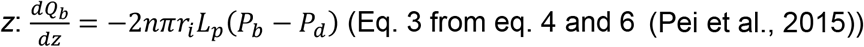

Change in dialysate flow rate with respect to

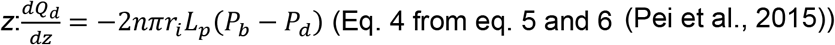

Change in blood and dialysate total bilirubin concentration multiplied by flow rate with respect to *z*: This equation is modified from eq. 7 and 8 (Pei et al., 2015). Unlike the previous model, we account for the change in the direction of convection across the dialyzer membrane when dialysate pressure exceeds blood pressure. A purely diffusive mode is introduced for small values of *J*_*v*_ to avoid division by zero errors when calculating *f* (see below).

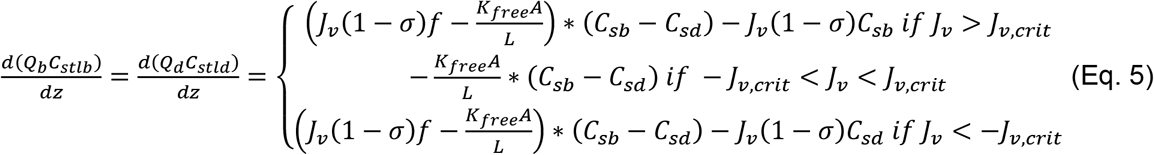

*J*_*v,crit*_ is calculated by the following criterion:

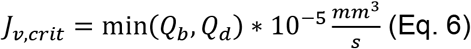

Blood free bilirubin in terms of blood total bilirubin:

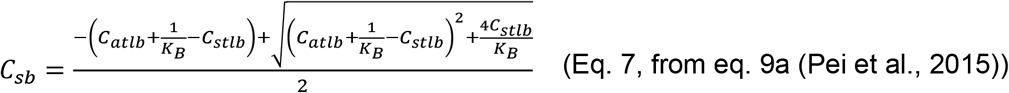

Dialysate free bilirubin in terms of dialysate total bilirubin:

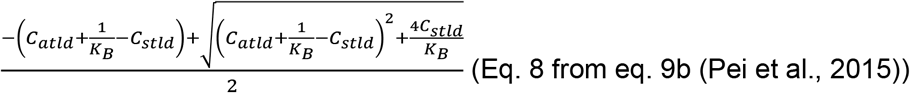

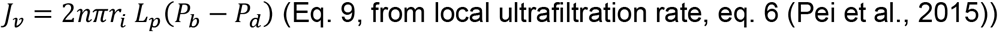

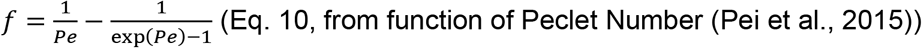

The equation for Peclet number is derived from eq. 9 (Villarroel et al., 1977)

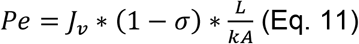

As flow rate changes, albumin concentration also changes, but since flow rate is periodic, so is albumin concentration. Thus, it doesn’t change over time, only spatially.

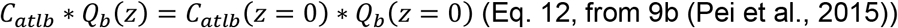

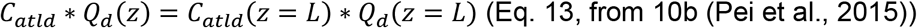

Time dependence is described as follows:

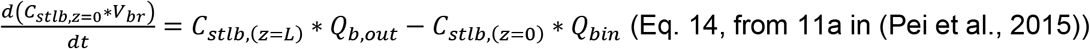

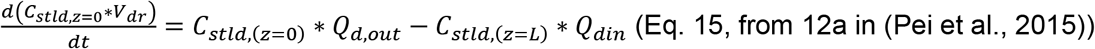

Because of numerical instability, a shooting method was used to solve the boundary value problem posed by counter-current flow in the dialyzer. The model was implemented using Julia Version 1.9.3 (Bezanson et al., 2017) with the packages Revise.jl, XLSX.jl, DifferentialEquations.jl (Rackauckas and Nie, 2017), FileIO, JLD2, ODEInterfaceDiffEq (Rackauckas and Nie, 2017), NonlinearSolve (Rackauckas et al., 2023), NLSolve (Mogensen et al., 2020), LinearAlgebra, and Roots (Verzani, 2020). We used the Julia function linspace provided by Jonathan Bieler (Bieler, 2020). We used the Julia remove function provided by Michael Franco (Franco, 2019). We used the myfind Julia function (“myfind,” 2017). The radau solver was used for the spatial solution (Hairer and Wanner, 1999). The built-in solver selection algorithm of DifferentialEquations.jl was used for the temporal differential equation solution (Rackauckas and Nie, 2019).

Details of the algorithm used to achieve numerical stability in solving this system are available in our provisional patent application (Novokhodko et al., 2024). Parameter sweeps were conducted on the University of Washington’s Hyak Supercomputer. Goodness of fit was analyzed by minimizing the sum of squares (goodness of fit to the entire solution) or by minimizing the percent error at the end of the trial (goodness of fit to the equilibrium bilirubin concentration). The two criteria are shown in Eq. 16 and Eq. 17 respectively. Here, *t* is time during the trial. *t*_*end*_ is the time at the end of the trial. *C*_*true*_ is the measured concentration. *C*_*model*_ is the concentration predicted by the model. *i* is an index that varies from 1 to *n* where *n* is the number of time points.

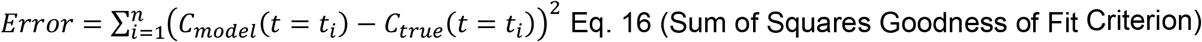

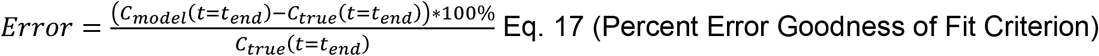

### 2.5 Experimental Procedure

Five sets of experiments were done in total:

- Condition 1: A pilot study (*n* = 3) using Bovine Serum Albumin (BSA) on both sides of an F6HPS dialyzer (Fresenius, Waltham, MA, USA) with blood and dialysate flow rates of *Q*_*b*_= 180 mL/min and *Q*_*d*_ = 90 mL/min. This was used to fit two parameters:

1. The dialyzer diffusive mass transfer coefficient for free bilirubin moving through polysulfone (*K*_*free*_*A*)
2. The bilirubin binding equilibrium constant for the primary binding site on BSA (*kB,BSA*)
  - Condition 2: A validation data set (*n* = 3) using BSA on both sides of an F3 dialyzer (Fresenius, Waltham, MA, USA) with blood and dialysate flow rates of 150 mL/min
  - Condition 3: A test data set (*n* = 3), like condition 2, but with the dialysate flow rate set to 20 mL/min, as previously described in a pilot study of a novel albumin dialysis system (Ahmad et al., 2024)
  - Condition 4: A test data set (*n* = 3), like condition 2, but with the dialysate flow rate set at 800 mL/min, which is the highest flow rate we have found in clinical practice (Leypoldt et al., 1997)
  - Condition 5: A test data set (*n* = 3), like condition 2, but with the dialysate flow rate set at 2 mL/min, which is relevant in some neonatal dialysis applications (Battista et al., 2023)

The precise values for the setups are summarized in Table 3. The F6HPS trials (setup 1) was used to set model parameters. These were the toxin binding affinity for albumin, and the dialyzer mass transfer-area coefficient for free bilirubin diffusion (*K*_*free*_*A*). The F3 trials were used to independently validate model predictions. The *K*_*free*_*A* value was adjusted between the two dialyzers (see Section 2.7). Flow rate was set using Masterflex Pumps with model numbers 07551-20 and 77521-40, using Easy-Load II Rotors with Model Number 772990-62 (Cole-Parmer, East Bunker, CT, USA) except for the 2 mL/min trial, in which an Ismatec 78017-10 (IDEX Corporation, Northbrook, IL, USA) pump was used on the dialysate side. Pressure was controlled using four RSCDRRE015PGSE3 pressure sensors (Honeywell International Inc, Charlotte, NC, USA) to prevent ultrafiltration, except in the 800 mL/min trial, where more robust PS04-G100KP-A4W pressure sensors (Same Sky, Lake Oswego, OR, USA) were used. Data from Honeywell pressure sensors were obtained using a Raspberry Pi 3B (Raspberry Pi, Cambridge, England, UK). Same Sky pressure sensor readings were processed using an HX711 (Sparkfun, Niwot, CO, USA) connected to an Arduino Uno (Arduino SA, Chiasso, Switzerland). Pressure was adjusted using c-clamps. The blood and dialysate reservoirs were immersed in a 37 °C water bath (Benchmark, Sayreville, NJ), except in Setup 1. There, temperature was measured by K type thermocouples connected to a Measurement Advantage DAQ and maintained near 37 °C on both sides using hotplates. Hotplates were from Corning (Glendale, AZ, USA), with model numbers PC-420, and PC-420D. K-type thermocouples were from MN Measurement Instruments (St. Paul, MN, USA). The DAQ was from Measurement Computing (Norton, MA, USA). Figure 1 shows diagrams of the circuits for each setup.

**Figure 1:**
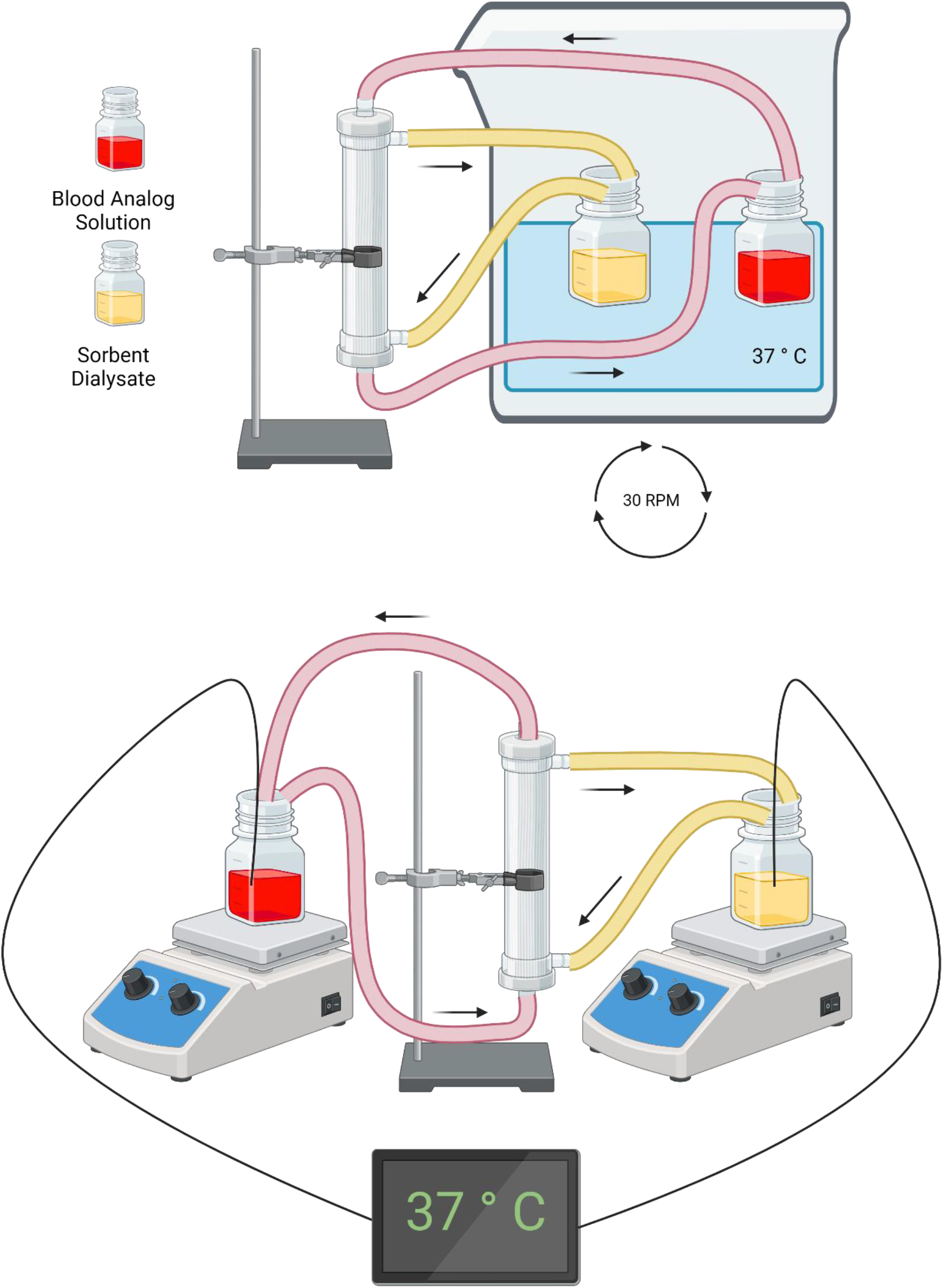
Experimental setup for conditions 2-5 (top) and condition 1 (bottom). Figure created with BioRender.

Solute concentrations, along with references supporting their clinical relevance, are listed in Table 2. pH on the blood and dialysate side was maintained between 7.35 and 7.45 using a pH Meter (OrionStar A211, Thermo Scientific, Waltham, MA), except in Setup 1, where the pH was maintained between 7.2 and 7.5 using the less precise pHoenix XL meter from MesaLabs. For Setup 1, pH in this was initially set at 7.2 ± 0.1 following previous work on protein bound toxin removal by our group (Pei et al., 2015). Acidosis is common in CKD patients (Marano et al., 2016; Noh et al., 2007). However, pH values below 7.3 are unusual. Acknowledging this, we increased the pH set point in later Setup 1 trials to 7.4 ± 0.1. In all setups, pH adjustment was done using 1 N HCl and NaOH. pH was set at 35-40 °C, determined using hotplates and thermocouples as previously described. In all setups care was taken to protect the setup from light to minimize photodegradation of bilirubin. For Setup 1, dialyzers were reused after being cleaned according to established clinical protocols (Cheung et al., 1999; Leypoldt, 2008). Bilirubin and Albumin measurement protocols were the same as in our previous work (Ahmad et al., 2024).

**Table 2:** Toxin concentrations in blood analog solution.

| Solute | Concentration |
| --- | --- |
| BSA | 2 g/dL (Bai et al., 2019) |
| Bilirubin | 20 mg/dL (Senf et al., 2004; Weiping, 2010) |
| Cholic Acid | 19.3 μM (Lewis et al., 1969) |
| Creatinine | 15 mg/dL (ul Amin et al., 2014) |
| Indoxyl Sulfate | 4 mg/dL (Barreto et al., 2009) |
| Copper | 21.56 μM (Rahelić et al., 2006) |
| Manganese | 2.5 μM (Rahelić et al., 2006) |

**Table 3:**
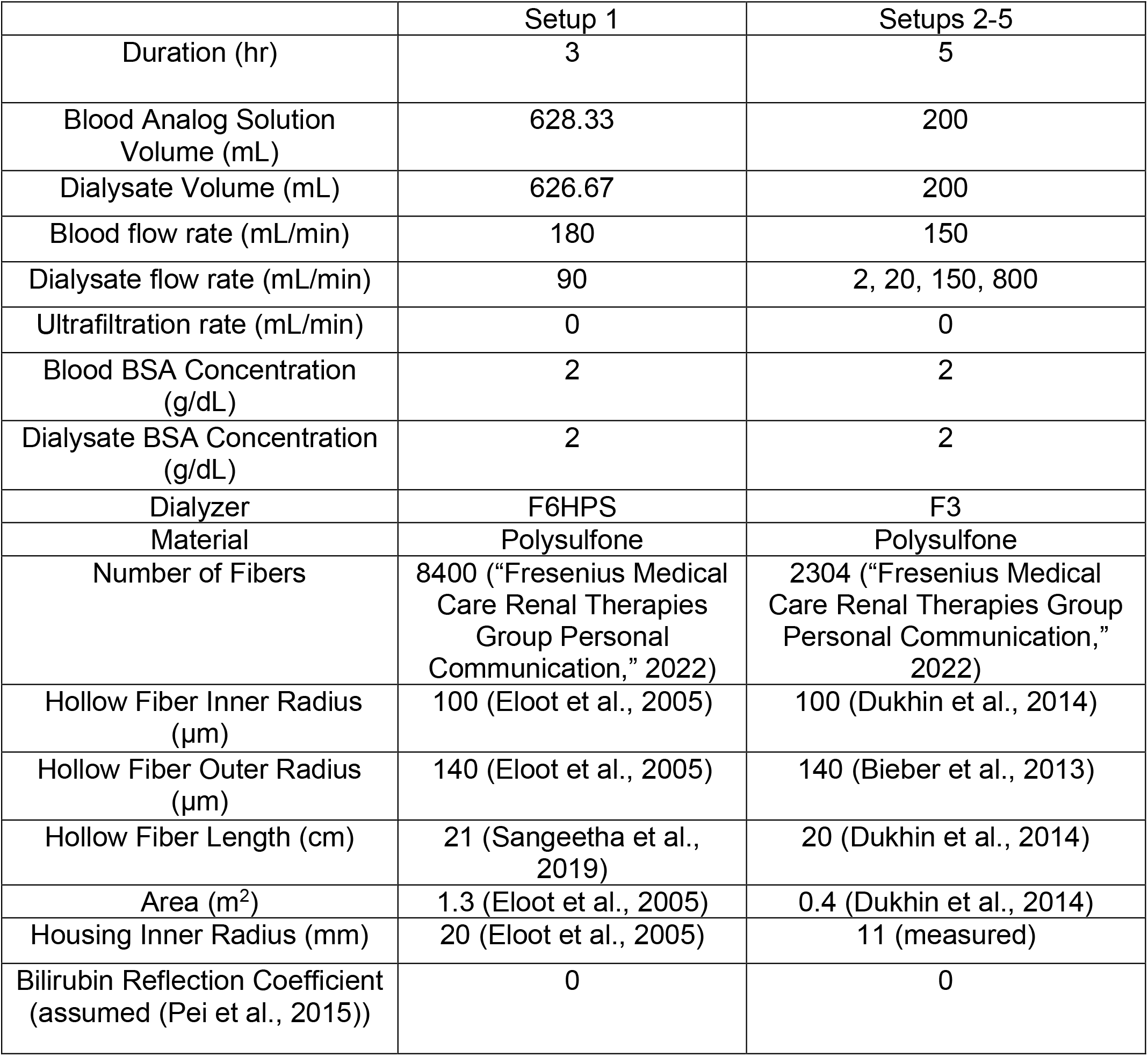
Albumin Dialysis Setups Used in This Study.

### 2.6 Polysulfone Membrane Hydraulic Permeability

Polysulfone membrane hydraulic permeability was measured by a method based previous studies (Liao et al., 2005), with some modification. A small dialyzer, called a “mini-module” was created out of individual F6HPS dialyzer fibers in a custom-made casing. Figure 2 shows the test setup. The formula for hydraulic permeability was:

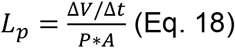

Δ*V* refers to the ultrafiltration volume. Δ*t* is the time. *A* is the total area of the dialysis membrane in the mini-module. Here, pressure is calculated as:

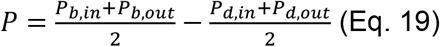

**Figure 2:**
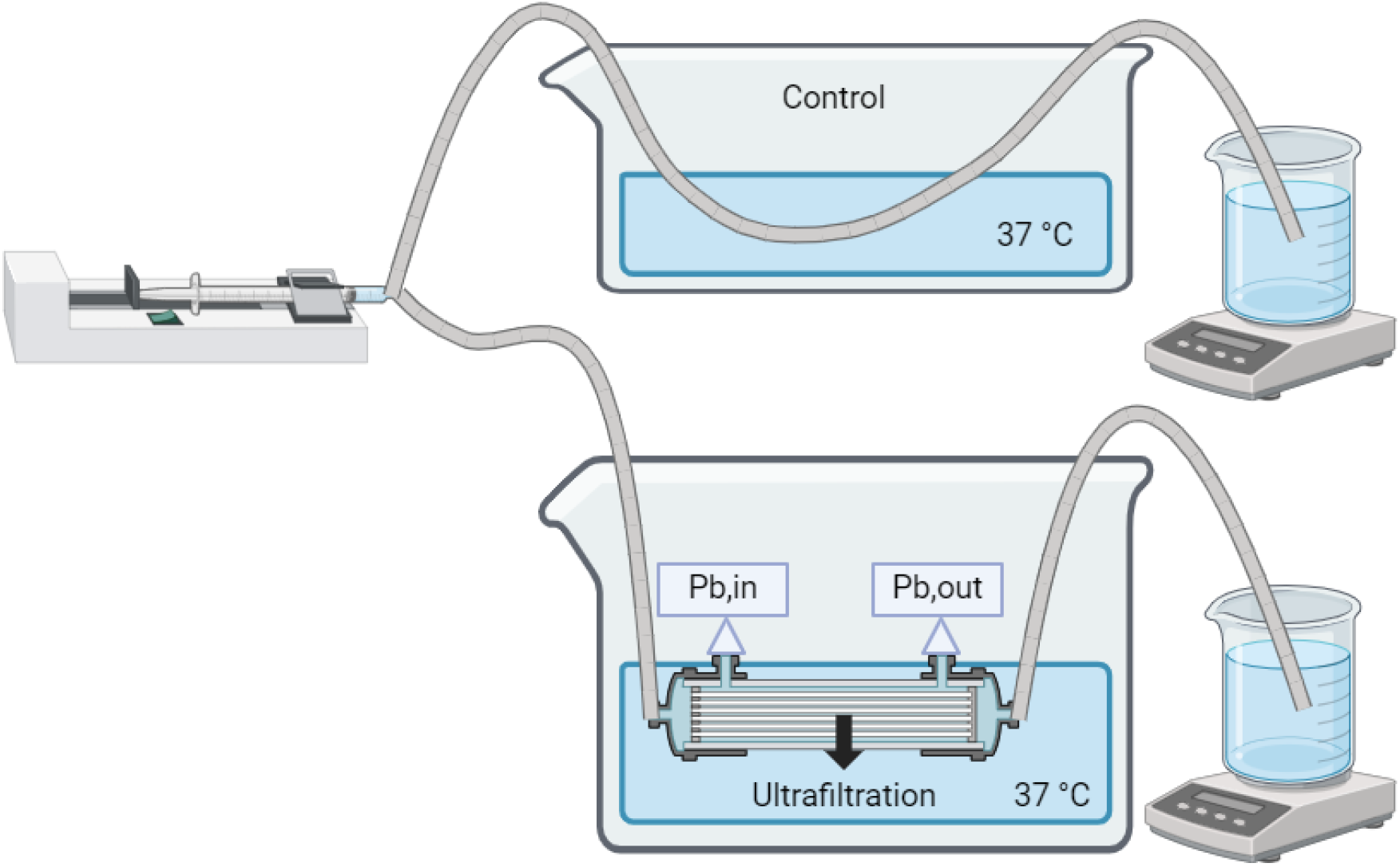
Hydraulic Permeability Test Setup. Figure created with BioRender.

In this formula, *P*_*b,in*_ is the inlet blood pressure and *P*_*b,out*_ is the outlet blood pressure. *P*_*d,in*_ and *P*_*d,out*_ are the corresponding dialysate pressures. Since the module is near the surface of a stirred water bath and the dialysate side is in contact with the water (Figure 2), the dialysate pressures are approximately equal and negligible.

Mini-modules were constructed out of F6HPS dialyzer polysulfone fibers. Fibers were carefully selected to be straight, to avoid any damage or kinking. Two mini-modules were constructed (designated F6HPS and F6HPS2). They were then bundled together in a fitting using Gorilla Epoxy (Gorilla Glue, Cincinnati, OH, USA). A neMESYS Low Pressure Syringe Pump (Cetoni, Korbussen, Germany) was used to set precise flow rates. Pressure on the inlet and outlet of the mini-module was measured using PX409-USBH sensors (Omega Engineering, Norwalk, CT, USA). Mini-modules were immersed in a 37 °C water bath. Heating, stirring, and temperature control were provided using hotplates, K-type thermocouples, and a data acquisition unit (DAQ). Hotplates were from Corning (Glendale, AZ, USA), with model numbers PC-420, and PC-420D. K-type thermocouples were from MN Measurement Instruments (St. Paul, MN, USA). The DAQ was from Measurement Computing (Norton, MA, USA). Flow through the mini-module was compared to flow through a control tube. The difference was the ultrafiltration rate Δ*V*. Following past work (Liao et al., 2005), flow rates of 1910 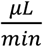, 764 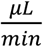, and 191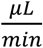 were tested for both mini-modules. One of the 191 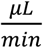 tests was an extreme outlier, so it was repeated, and 382 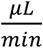 was also tested for that mini-module. These additional tests did not replicate the outlier value, so the outlier value was then discarded.

### 2.7 Fit Parameters

Two parameters were fit computationally rather than being determined from the literature. The first was the primary binding constant for bilirubin-BSA binding. All literature values for this parameter fall within a narrow range (Table 4). The primary binding site of bilirubin on BSA has approximately an order of magnitude greater affinity than the secondary binding site (Chen, 1973). Thus, the secondary binding site was neglected for this analysis. the constant for the primary binding site was fit. The lower bound of the search space was set to 0.5E7 (1/M). The upper bound was set to 7.5E7 (1/M). The space was searched in steps of 1E7 (1/M).

**Table 4:**
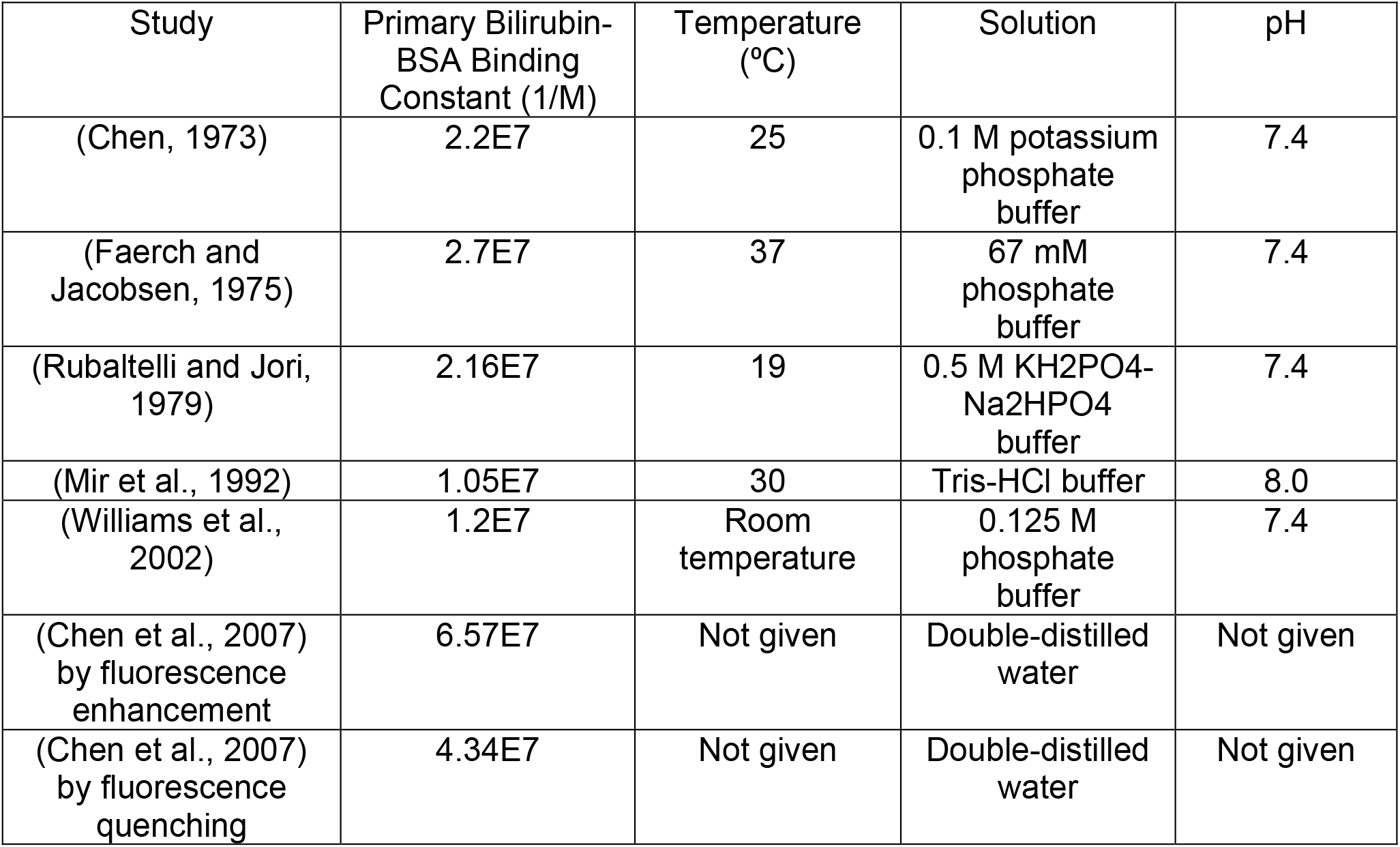
Reported values of bilirubin-BSA binding constant.

The second fit parameter was the product of the dialyzer mass transfer-area coefficient for free bilirubin diffusion and area (*K*_*free*_*A*). The highest *KoA* value reported in the literature is 2000 mL/min (Meyer et al., 2007). Thus, this parameter was optimized over the range from 100 to 2500 mL/min in steps of 100 mL/min. In Equation 5 this coefficient is analogous to *KoA* for toxins which are not protein bound, but there are important differences. This coefficient does not relate directly to the clearance of the toxin of interest by the standard relationship (Michaels, 1966). This calculation assumes that the mean concentration difference in counter-current dialysis is the logarithmic mean of the inlet and outlet concentration differences. For free bilirubin, this is not the case. On the blood side, as free bilirubin crosses the membrane the unbound concentration declines. This creates a thermodynamic driving force for the release of additional bilirubin from albumin. In contrast, on the dialysate side as bilirubin crosses the membrane, it binds to free binding sites on dialysate albumin. This lowers the free dialysate side bilirubin concentration. Additionally, this coefficient depends on the assumption that blood and dialysate side clearance are approximately equal. Other processes such as membrane binding of bilirubin may cause blood side clearance to exceed dialysate side clearance. Polysulfone membranes have been observed to bind small quantities of bilirubin (Peng et al., 2016). This would lead to an apparently elevated *K*_*free*_*A* being the best fit parameter because it must account for other processes such as membrane binding.

Afterwards, *K*_*free*_*A* was adjusted to account for a new dialyzer following equation 20. Because the F3 and F6HPS dialyzers are both polysulfone, it was assumed that area was the only adjustment needed, meaning the two dialyzers had the same value of *K*_*free*_. The dependence of *K*_*free*_*A* flow rate (Ouseph and Ward, 2001) was neglected.

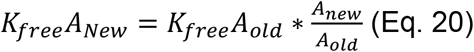

## 3 Results

### 3.1 Hydraulic Permeability

Hydraulic permeability was measured as 8.61*10^−11^ m/(s*Pa) ± 2.14*10^−11^ m/(s*Pa) (average ± standard deviation). This measurement was used for subsequent modeling. This is much lower than the value obtained by Liao and colleagues for polysulfone of 1.628*10^−9^ m/(s*Pa) (Liao et al., 2005). However, it is comparable to the value obtained by Benavente and Jonsson of 10^−10^ m/(s*Pa) (Benavente and Jonsson, 2000). Hydraulic permeability test raw data is reported in Table 5. The discarded outlier test is highlighted in bold. Averages and standard deviations are reported in Table 6.

**Table 5:** Hydraulic Permeability Measurements from Individual Trials. *Lp* refers to hydraulic permeability. *P*_*b,in*_ refers to blood side (luminal) inlet pressure. *P*_*b,out*_ refers to blood side outlet pressure. *ΔV* refers to the change in volume. The test highlighted in bold was an outlier which was discarded.

| $L_p$<br>(m/s*Pa) | Mini-Module | Flow Rate<br>(μL/min) | $P_{b,in}$ (cm<br>H<sub>2</sub>O) | $P_{b,out}$ (cm<br>H<sub>2</sub>O) | $\Delta V$ (mL) | Time<br>(min) |
| --- | --- | --- | --- | --- | --- | --- |
| 1.013E-10 | F6HPS2 | 1910 | 209.7 | 3.94 | 0.628 | 10.5 |
| 1.020E-10 | F6HPS2 | 764 | 65.139 | 2.615 | 0.383 | 20 |
| 9.601E-11 | F6HPS2 | 382 | 35.828 | 2.58 | 0.409 | 40 |
| **2.768E-11** | **F6HPS2** | **191** | **19.76** | **4.162** | **0.11** | **60** |
| 8.851E-11 | F6HPS2 | 191 | 21.127 | 2.904 | 0.473 | 80 |
| 9.750E-11 | F6HPS | 1910 | 531.57 | 3.726 | 1.45 | 10 |
| 7.463E-11 | F6HPS | 764 | 269.35 | 1.955 | 1.122 | 20 |
| 4.257E-11 | F6HPS | 191 | 50.8 | 4.507 | 0.522 | 80 |

**Table 6:** Average and standard deviation hydraulic permeability values for both mini-modules and overall.

| Mini-Module | Average $L_p$ (m/s*Pa) | Standard Deviation (m/s*Pa) |
| --- | --- | --- |
| F6HPS | 7.157E-11 | 2.75928E-11 |
| F6HPS2 | 9.696E-11 | 6.233E-12 |
| Total | 8.61E-11 | 2.14E-11 |

### 3.2 Albumin Dialysis Study

Table 7 summarizes measured average starting bilirubin and albumin concentrations for all five setups. These values were used for modeling to avoid error caused by variations in the initial solution composition.

**Table 7:** Average Initial Bilirubin and Albumin Concentration. *N* = 3 for all conditions. Values are shown as mean ± standard deviation.

| Condition | Starting Blood<br>Bilirubin (mg/dL) | Starting Blood<br>Albumin (g/dL) | Starting Dialysate<br>Albumin (g/dL) |
| --- | --- | --- | --- |
| 1 (F6HPS, $Q_b = 180$<br>mL/min, $Q_d = 90$<br>mL/min) | $15.37 \pm 3.06$ | $2.24 \pm 0.37$ | $1.77 \pm 0.41$ |
| 2 (F3, $Q_b = Q_d = 150$<br>mL/min) | $18.81 \pm 2.84$ | $1.90 \pm 0.07$ | $1.94 \pm 0.23$ |
| 3 (F3, $Q_b = 150$<br>mL/min, $Q_d = 20$<br>mL/min) | $16.76 \pm 2.35$ | $2.13 \pm 0.12$ | $2.29 \pm 0.13$ |
| 4 (F3, $Q_b = 150$<br>mL/min, $Q_d = 800$<br>mL/min) | $18.28 \pm 0.43$ | $2.03 \pm 0.02$ | $2.10 \pm 0.04$ |
| 5 (F3, $Q_b = 150$<br>mL/min, $Q_d = 2$<br>mL/min) | $17.36 \pm 1.94$ | $2.15 \pm 0.22$ | $2.21 \pm 0.14$ |

### 3.3 kA and kB Parameter Sweep

The optimal values of *kB,BSA* and *K*_*free*_*A* for bilirubin removal Setup 1 were found to be *kB,BSA* = 0.5E7 (1/M) and *K*_*free*_*A* = 2500 mL/min. This result was not sensitive to changing the number of fibers from *n* = 8400 provided by Fresenius to *n* = 9200 (Eloot et al., 2005). The results are shown in Figure 3. Panel A shows the parameter sweep results for the sum of squares criterion. Panel B shows the parameter sweep results for the percent error criterion. Figure 4 shows the model prediction compared to the true measurements.

**Figure 3:**
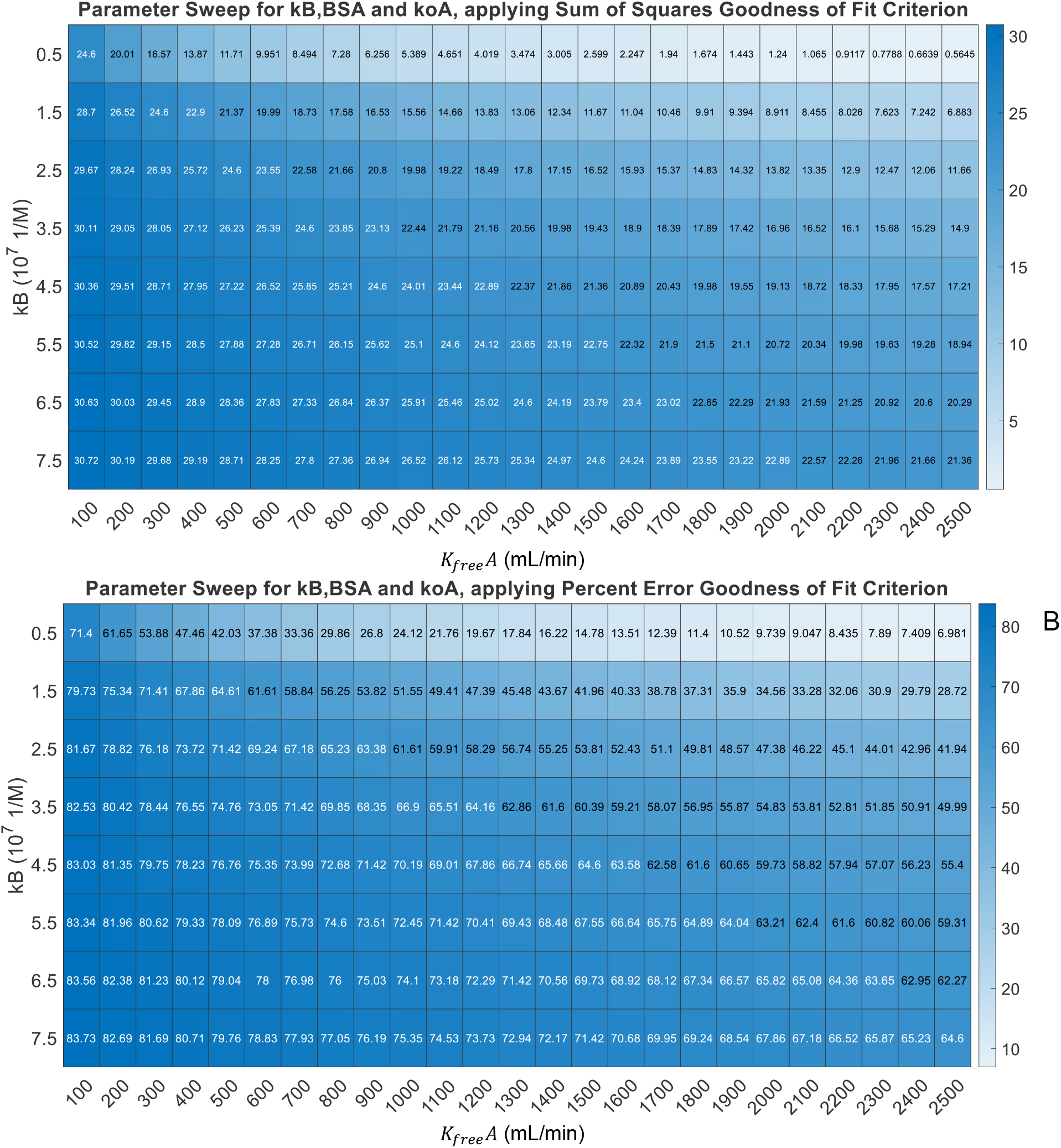
A: Parameter sweep results for *kB,BSA* for bilirubin and *K*_*free*_*A*, measuring best fit by the sum of squares criterion. B: Parameter sweep results for *kB,BSA* for bilirubin and *K*_*free*_*A*, measuring best fit by the percent error criterion.

**Figure 4:**
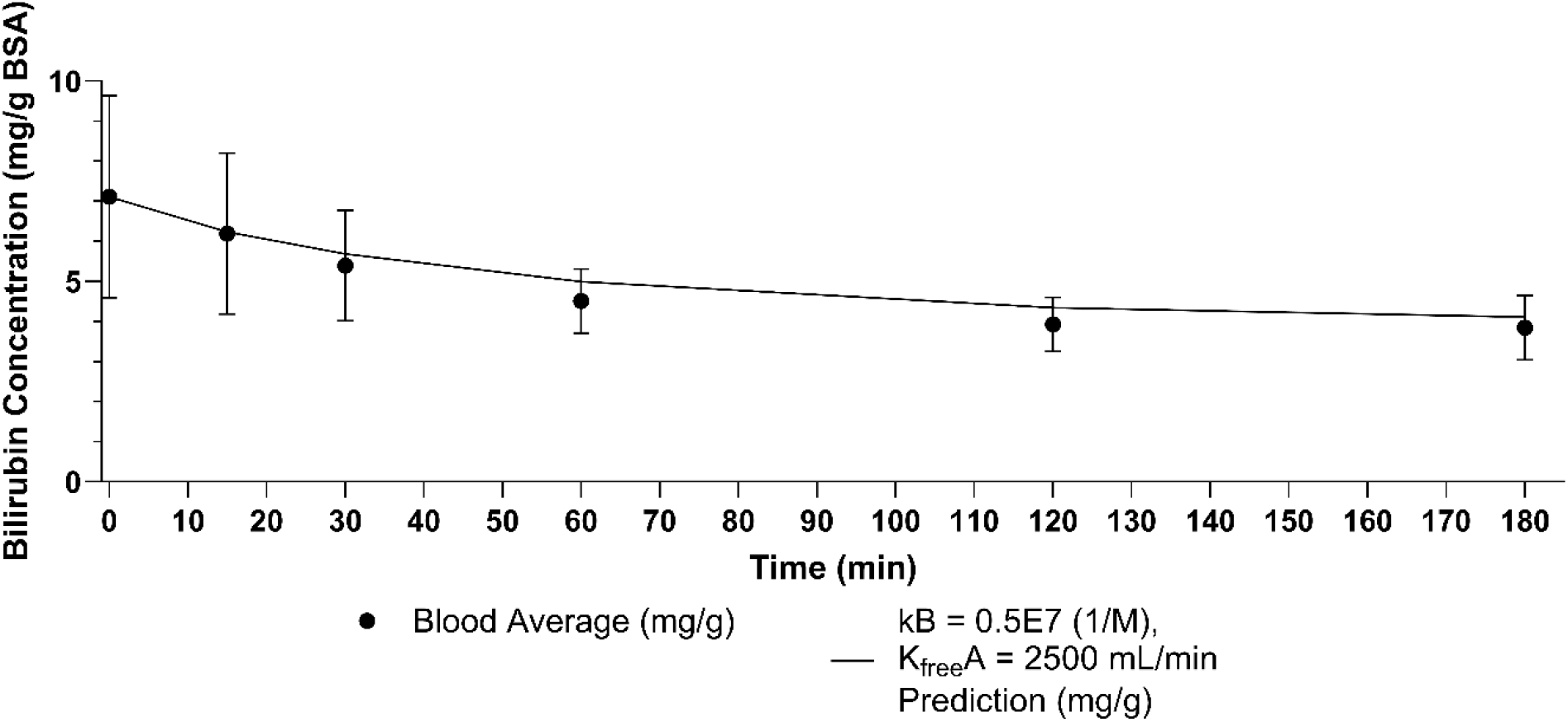
Blood side reservoir bilirubin over time for F6HPS with *kB* = 0.5E7 (1/M) and *K*_*free*_*A* = 2500 mL/min and *n* = 8400 compared to experimental data. Error bars are standard deviation. Final percent error 6.98%. Sum of squares error 0.56.

### 3.4 Model Validation for F3 Dialyzer and Flow Rates

Next, we validated the model using Condition 2, then tested it against the other three flow rates for the F3 polysulfone dialyzer. Applying Equation 20, the new *K*_*free*_*A* for this dialyzer was 770 mL/min. The results of the simulation and experiments are summarized in Figure 5. Experimental results are summarized in Figure 6. Individual conditions are shown in Figure 7.

**Figure 5:**
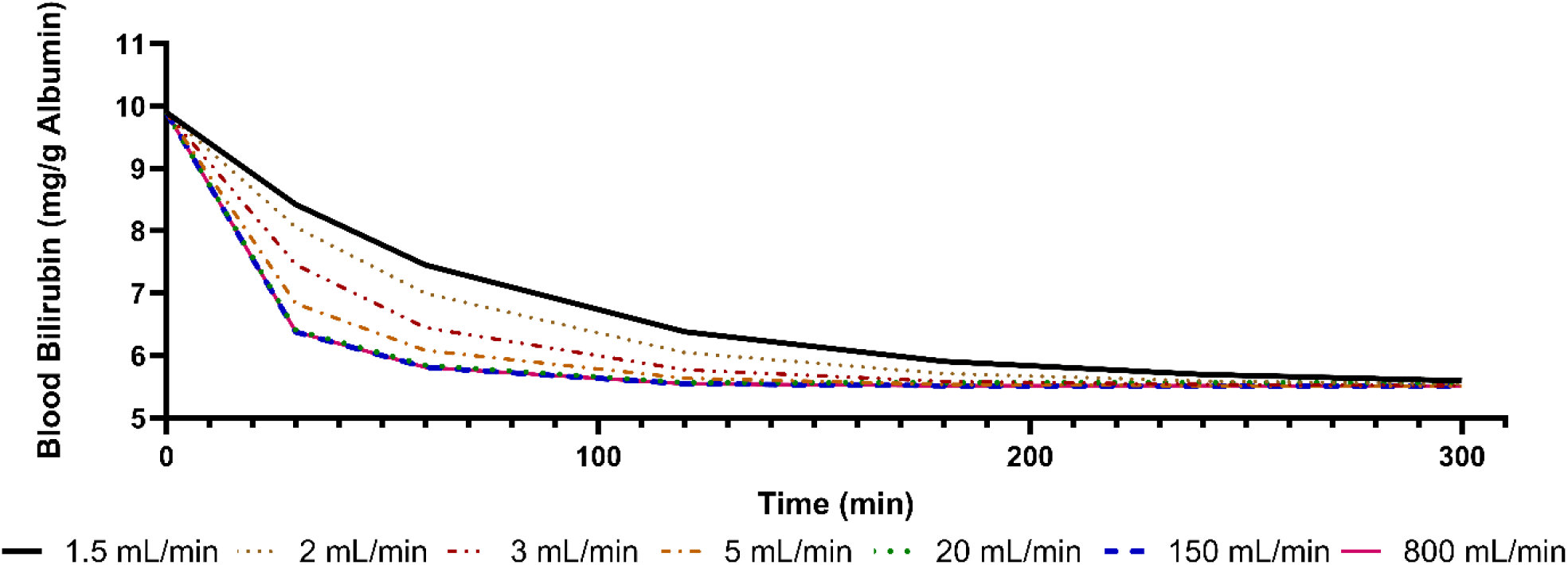
Model predictions for bilirubin removal with an initial blood side bilirubin concentration of 20 mg/dL, and blood and dialysate side BSA concentrations of 2 g/dL. Each line corresponds to a dialysate side flow rate. The 20 mL/min, 150 mL/min and 800 mL/min lines overlap.

**Figure 6:**
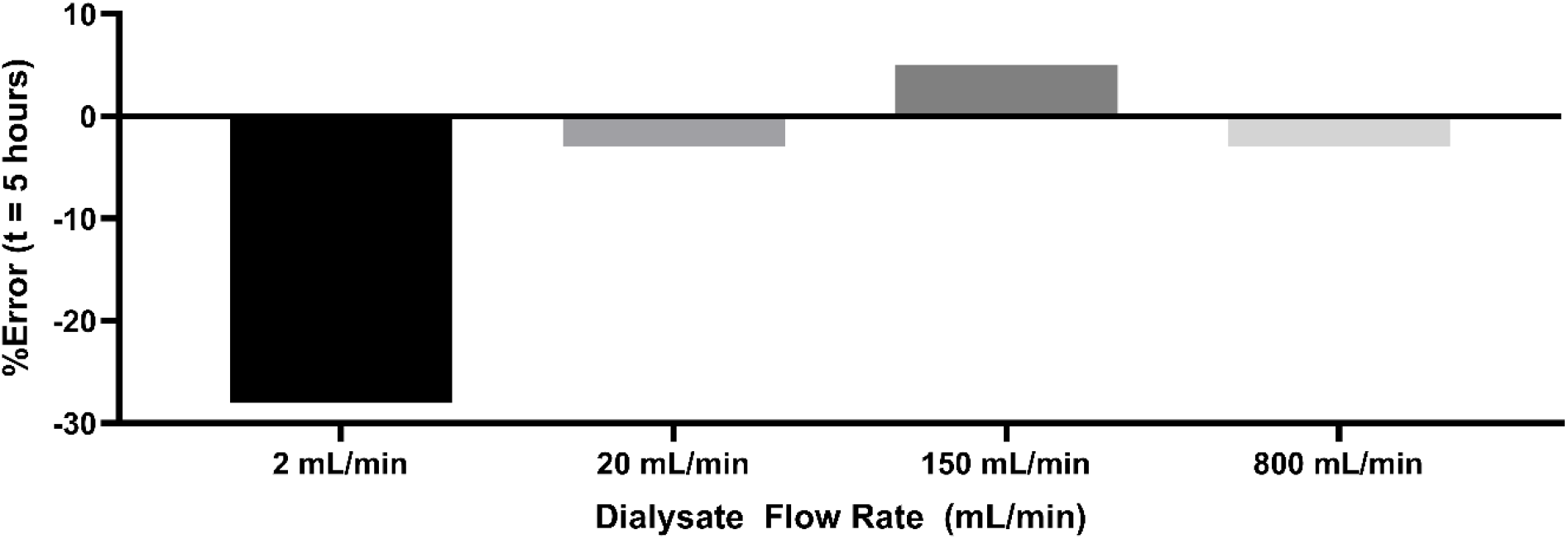
Errors for final bilirubin concentration for F3 dialyzer setup with various flow rates. Numerical values given in Table 8.

**Figure 7:**
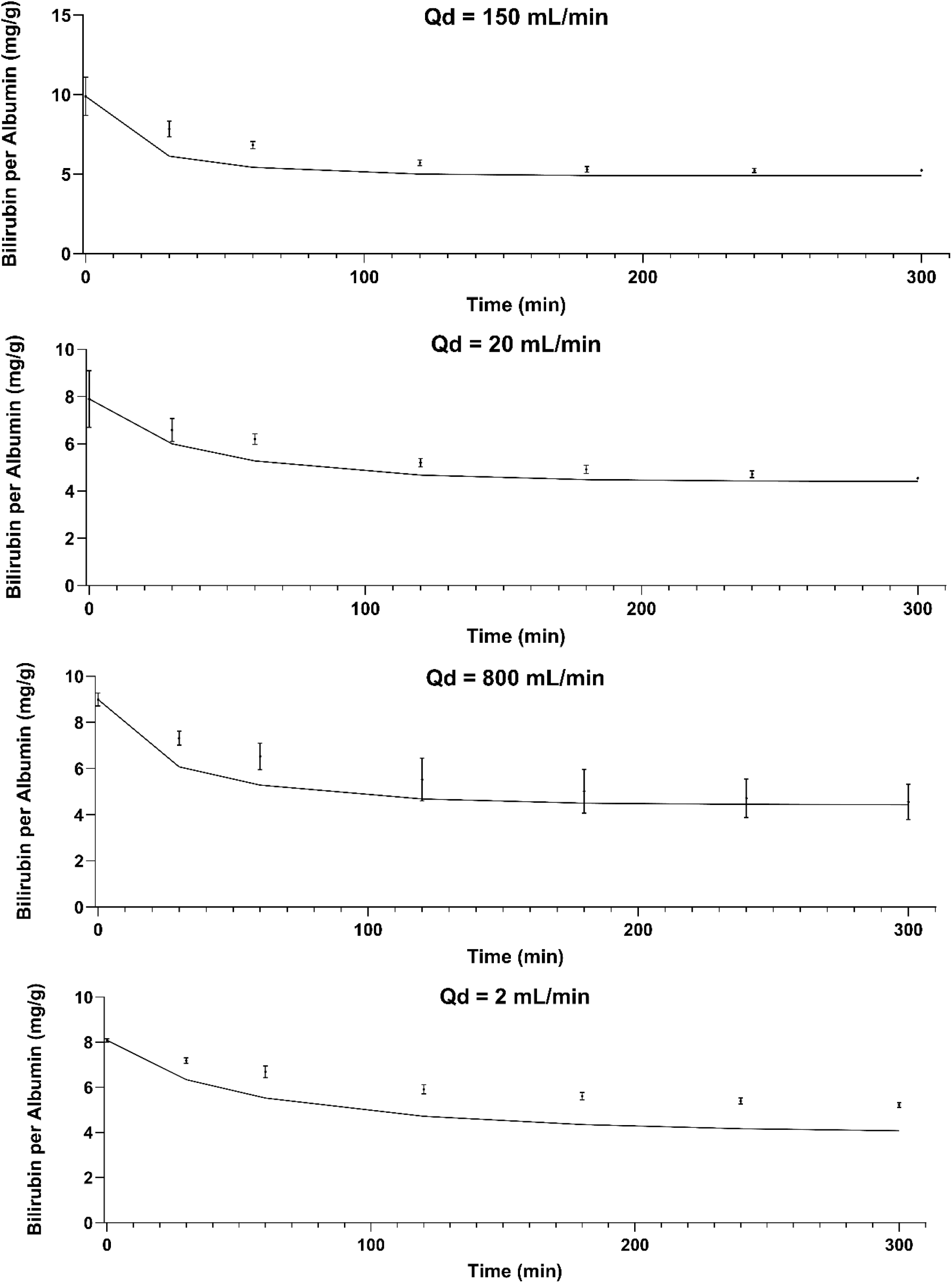
Results and model predictions for dialysate flow rates of 150 mL/min (Panel A), 20 mL/min (Panel B), 800 mL/min (Panel C), and 2 mL/min (Panel D). Errors bars are standard deviation. Where error bars are not shown, it is because they would be smaller than the data point depicted on the graph. Points are experimental data and the line is the model prediction.

Figure 5 shows the model predictions for flow rates ranging from 800 mL/min to 1.5 mL/min with a starting bilirubin concentration of 20 mg/dL and starting albumin concentrations on the blood and dialysate side of 2 g/dL. Figure 6 summarizes the % error of the final concentration for all flow rates. Figure 7 shows results for individual conditions and how they compare to model predictions. Table 8 summarizes the accuracy of the model in predicting the outcomes in these conditions. For all flow rates measured except 2 mL/min, the model predicted the final blood bilirubin concentration within 7%.

**Table 8:**
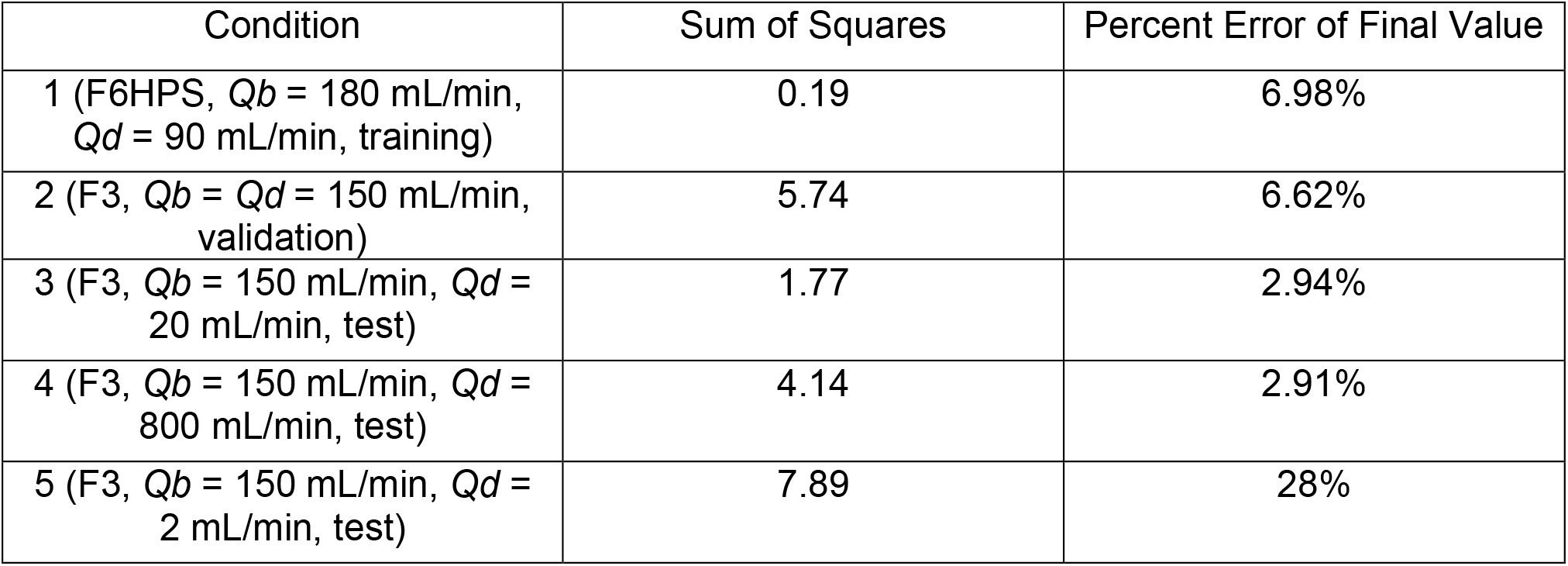
Goodness of Fit for Different Conditions by Sum of Squares and Percent Error of Final Value Criteria.

## 4 Discussion

This work validates a thermodynamically-based computational model that can be used to rationally design albumin dialysis. This paves the way for optimization of an albumin dialysis system for protein bound toxin removal. It also creates the potential for personalized dialysis regiments, where patients are given flow rate conditions and dialysate compositions tailored to their body volume and toxin concentrations. This would play a similar role to the NxStage Dosing Calculator in home dialysis (Molina et al., 2018). This work will guide us in optimizing AMOR treatment prescriptions.

This work only analyzed the model’s ability to predict the impact of changes in dialyzer and dialysate flow rate. Further work is in progress to predict the impact of changes in blood flow rate and other parameters. Another limitation was small scale. In the test conditions used to validate the model, blood volume was 200 mL – 630 mL. In a patient the plasma volume is approximately 3 L (Sharma and Sharma, 2024). A larger scale *in vitro* test is needed to validate the model’s predictive ability in patients.

The model correctly predicts the equilibrium outcomes of albumin dialysis, but it overestimates the kinetics of bilirubin transport across the membrane. This appears mathematically in an implausibly high *K*_*free*_*A* value. For example, it predicts a *K*_*free*_*A* value for bilirubin of 2500 mL/min for the F6HPS dialyzer, whose *kA* for urea is only 746 mL/min under similar flow conditions (Li et al., 2010). Pei and colleagues previously reported a bilirubin *kA* value of 800 mL/min for a Gambro 6LR dialyzer using a similar model (Pei et al., 2015). The *KoA* for urea for the Gambro 6LR dialyzer is 736 mL/min (Ahmad et al., 2015). We could not replicate their result with a 800 mL/min *kA* using their model. A higher *kA* value would be needed. In their original work on protein bound toxin removal modeling, Patzer and Bane noted that the dialyzer mass transfer coefficient of their membranes increased after 180 minutes (3 hours) of dialysis (Patzer and Bane, 2003). They suggested that bilirubin and albumin binding to membrane pores accounts for this phenomenon. It would be very useful to accurately predict bilirubin kinetics in our model so that it can be coupled with models of bilirubin absorption onto activated charcoal to predict the behavior of a combined dialysis/absorption system like MARS. Thus, we will extend the model to incorporate bilirubin binding to albumin absorbed onto the dialysis membrane.

## 5 Conclusion

We present a computational model that uses the fundamental physics of flow rate, pressure, and concentration in the dialysis process to predict the rate of protein bound toxin removal using albumin dialysate. We validate this model against experimental data at four different flow rates and for two different polysulfone dialyzers. For all conditions except the lowest flow rate (2 mL/min), the model produces low-error predictions. Final values are predicted with less than 7% error. This demonstrates the value of this model for designing optimal albumin dialysis protocols.

## Supporting information

Figure 1 part 2 Publication License

Figure 2 Publication License

Figure 1 part 1 Publication License

## 6 Acknowledgements

This work was facilitated through the use of advanced computational, storage, and networking infrastructure provided by the Hyak supercomputer system and funded by the STF at the University of Washington.

Alexander Novokhodko acknowledges funding from the National Science Foundation Graduate Research Fellowship Program under Grant No. DGE-1762114. Any opinions, findings, and conclusions or recommendations expressed in this material are those of the author(s) and do not necessarily reflect the views of the National Science Foundation. Alexander Novokhodko acknowledges funding from the Dempsey Startup Competition, Hollomon Health Innovation Challenge, and the Ron and Wanda Crockett Endowed Fellowship in Mechanical Engineering.

